# Polygenic Prediction of Complex Traits with Iterative Screen Regression Models

**DOI:** 10.1101/2020.11.29.402180

**Authors:** Meng Luo, Shiliang Gu

## Abstract

Although genome-wide association studies have successfully identified thousands of markers associated with various complex traits and diseases, our ability to predict such phenotypes remains limited. A perhaps ignored explanation lies in the limitations of the genetic models and statistical techniques commonly used in association studies. However, using genotype data for individuals to perform accurate genetic prediction of complex traits can promote genomic selection in animal and plant breeding and can lead to the development of personalized medicine in humans. Because most complex traits have a polygenic architecture, accurate genetic prediction often requires modeling genetic variants together via polygenic methods. Here, we also utilize our proposed polygenic methods, which refer to as the iterative screen regression model (ISR) for genome prediction. We compared ISR with several commonly used prediction methods with simulations. We further applied ISR to predicting 15 traits, including the five species of cattle, rice, wheat, maize, and mice. The results of the study indicate that the ISR method performs well than several commonly used polygenic methods and stability.

## Introduction

The continuous accumulation of genetic data in existing association analysis studies has led to increasing interest use of genetic markers to predict complex trait phenotypes and diseases^1-3^. In animals or plants, accurate phenotypic prediction using genetic markers can assist in selecting individuals that meet the needs of products (high breeding value) and can effectively promote breeding programs^4-6^. In human genetics, which accurate use of genetic markers for phenotypic prediction, especially the heritable and highly polygenic, can promote disease prevention and intervention^7,8^, such as, polygenic risk scores that have shown promise in predicting human complex traits and diseases, and may facilitate early detection, risk stratification, and prevention of common complex diseases in healthcare settings^8-12^. And the genotype information can be used to develop individualized drug delivery for customized treatment and predict possible outcomes^13^. In animals, such as cattle, producers have accepted the use of whole-genome selection techniques to evaluate and select offspring^14^. Besides, it also benefits plants. In wheat and maize, studies have shown that multi-cycle whole-genome selection can achieve better and desirable results^2,15-17^. Therefore, in recent years, researchers have regarded phenotype prediction as a critical step in joint functional genomics and genome-wide research^10,18^.

However, with the growth of high-throughput genomics data, accurate phenotype prediction requires the development of statistical methods that can simulate all or majors SNPs simultaneously^9,19,20^. Moreover, previous genome-wide association analysis studies have shown that many complex trait phenotypes and diseases have a polygenic genetic background, mainly controlled by many genetic variation sites with smaller effects. For example, in human genetics, Hundreds of mutation sites have been evaluated to affect human height and body mass index (BMI) ^21,22^, making the height and BMI of different groups of people diversified. Similarly, in the complex traits of animals and plants, there are phenotypic variations controlled by dozens of variation sites, such as traits related to rice yield composition^23^; features about cattle, such as back fat, milk yield, And carcass weight^24,25^. Because complex traits and common diseases have a multi-gene structure, only a few identified related mutation sites (SNPs) explain a small part of the phenotypic variation, so accurate phenotype and disease risk prediction cannot be drawn. On the contrary, accurate phenotype prediction requires a multi-gene model to be able to utilize all or major genome-wide SNPs genetic marker variations that explain the phenotype. In the past ten years, multi-gene models have been successfully developed and applied for prediction, and many animal breeding programs have been changed in the context of selection^26-29^. In addition, recently, the application of polygenic models in human GWASs has also achieved promised results^30-33^.

Most of the existing polygenic models used for prediction make assumptions about the distribution of effect sizes. The different methods are mainly due to the differences in the assumptions of these other models. For example, the commonly used linear mixed model (LMM), also known as the genomic best linear unbiased prediction (GBLUP)^34^, and rrBLUP that one of the first methods proposed for genomic selection was ridge regression (RR) which is equivalent to best linear unbiased prediction (BLUP) when the genetic covariance between lines is proportional to their similarity in genotype space^35^. And both assume that the size of the effect obeys a normal distribution^33,35^; also, the Bayes alphabetic included BayesA, and BayesB methods assume it the distribution of the effect size follows the t distribution or other distributions^36,37^; the effect size assumed by BayesC is also a normal distribution^36^; Bayes LASSO follows the double exponential and Laplace distribution^38,39^; the BSLMM assumption follows A mixture of two normal distributions^28^; while BayesR assumes a three-component normal distribution mixture^40^; Bayes no-parameter model (DPR, Dirichlet process regression)^19^ does not rely on any specific assumptions, but according to the Dirichlet Process Regression to give the hypothesis of a particularly suitable model. Given many model choices, people naturally think of which method can be used for any particular trait. Previous studies have shown that accurate prediction needs to choose a priori effect size distribution, which can be near consistent with the true effect size distribution. The inferred posterior can be well approximated to the traits with a multi-gene structure under consideration^30,40^. However, the priority of the effect size distribution for any particular trait or disease is unknown. Therefore, in order to maximize the model’s strong performance, the most important thing is to have a reasonable effect size distribution assumption, not the prior distribution while is flexible enough. As close as possible to the true effect distribution^28,40^.

For highly polygenic traits, it is assumed that the normal distribution can well fit the true effect size distribution. Therefore, LMM (linear mixed model) can obtain high predictive power^28,40^. As we all know, the effect size of each SNP site that causes phenotypic variation that can be divided into small effects, medium effects, and large effects (directly influence (or perfectly tag a variant that directly influences) the trait of interest, associated) which inferred that weak effect (small and medium) and strong effect ^27,41,42^; These classifications are based on ordinary least squares (OLS) effect size estimates for each SNP in a regression framework. The remaining loci have no effect (have no effect on the trait at all, non-associated). So if exiting a model make it true which is good enough to have identified all loci, can put all the loci are identified, and make use of these variable loci are also very reasonable to predict and prediction the result is a very good performance, such as, BayesR^28,40^. Here, we proposed the Iterative Screen Regression (ISR) also assumes that its effect size fits a normal distribution. In this study, the proposed Iterative Screen Regression model was used to explore the phenotype prediction and compared it with other commonly used methods in simulation and real phenotype prediction. We use simulation and real data applications to explain and analyze the advantages and disadvantages of ISR for phenotypic prediction. Results from ISR are compared with commonly polygenic prediction models, which included DPR, BayesR, BSLMM (Bayesian sparse linear mixed model), Bayes, BayesB, BayesC, BayesLASSO and rrBLUP and the genomic selection of 15 traits of five species and 10 complex traits of white mice will be used for genetic prediction analysis.

## Results

### Method overview

An overview of our method is provided in the Methods section. For details please see ISR^42^. Briefly, we offered a new regression statistics method and combined a unique variable screening procedure (Fig.1).

**Fig. 1.**
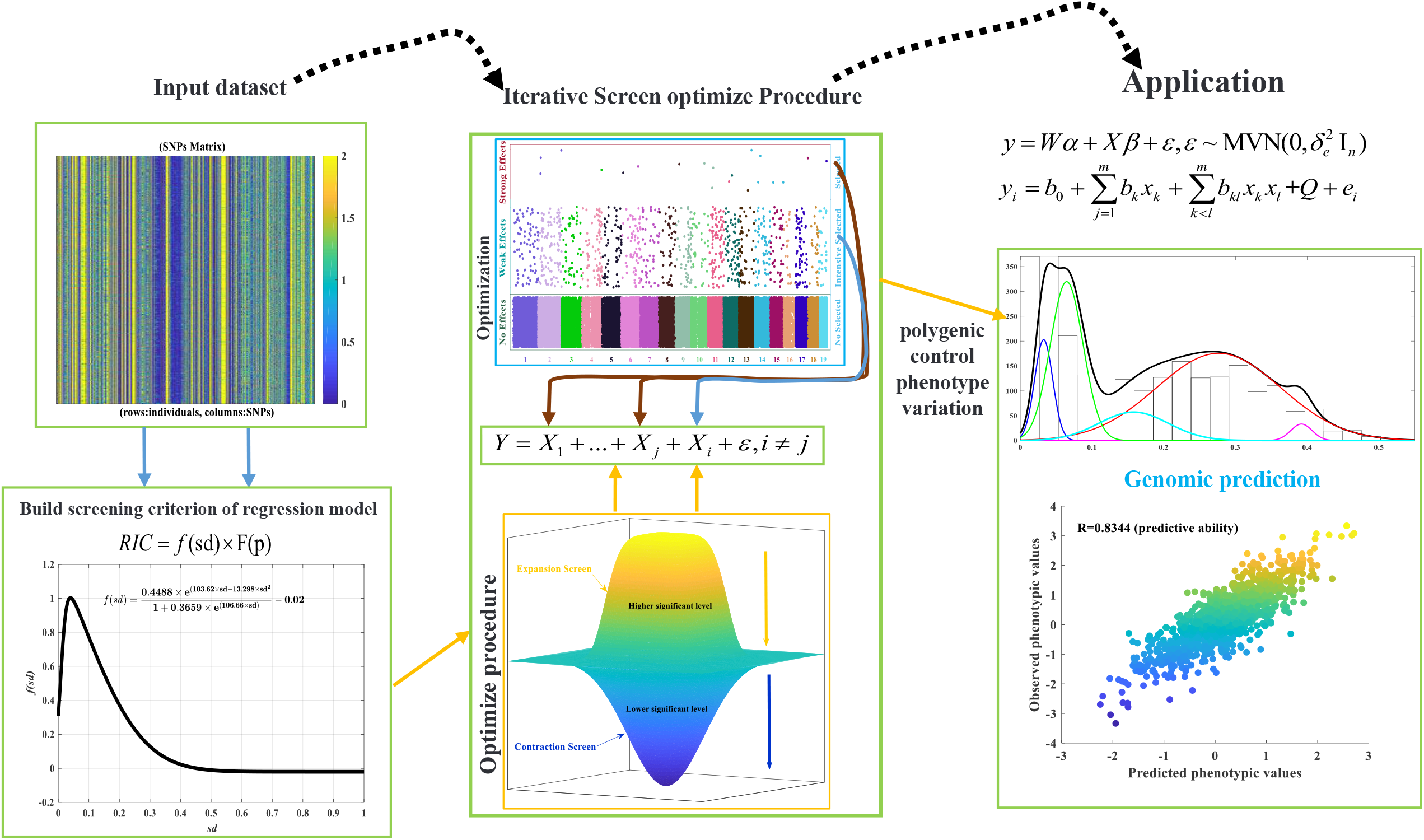
Schematic overview of model-based is iterative screening regression for GS. The first input dataset with markers (SNPs) matrix representing individual genotypes (rows) of a population with alleles (0, 2, and 1, missing genotypes will be replaced by the mean genotype or imputed by others complicate algorithm) per marker (columns). Secondly, we formulated a regression information criterion (RIC, objective function) as the screening criterion of the regression model. Combined the proposed iterative screen optimize the procedure, which mainly included expansion screen and contraction select two-steps. The third, apply it to multiple regression analysis, and two models can be selected, one for the linear model and the other for is the binomial model (including the epistasis effect). Here, we show the polygenic prediction of complex traits which the PHB phenotype distribution, where according to the character numerical simulation and we found the optimal equation that is five normally distributed superpositions and the black curve is explanation all models. Each of the models is blue curve, green curve, red curve, cyan curve, and purper curve (five major genes), and the best fitting model is finally selected as follows, and the optimal parameters estimated see the supplementary Table 6, From R2 =0.9982 (determination coefficient), it can be seen that the fitting degree is very high. This model can well explain the character (Figure 2). Except for b13 and b14, all the other T-tests reached a significant level.

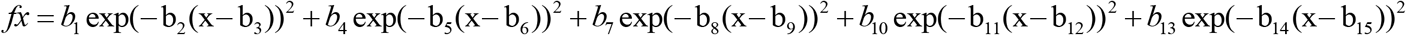

### Simulations

We first compare the performance of ISR with several other commonly used prediction methods using simulations. A total of seven different methods are included for comparison: (1) DPR; (2) BSLMM (GEMMA); (3) BayesA; (4) BayesB; (5) BayesC; (6) BayesLASSO; (7) rrBLUP. Note that DPR has been recently demonstrated to outperform a range of existing prediction methods (e.g., BayesR and MultiBLUP); thus, we do not include other prediction methods into comparison for polygenic prediction.

To make our simulations as real as possible, we used genotypes from an existing cattle GWAS dataset with 5024 individuals and 42,551 SNPs and simulated phenotypes. To cover a range of possible genetic architectures, we consider sixteen simulation settings from four different simulation scenarios with the phenotypic variance explained (PVE) by all SNPs being either 0.2, 0.5, or 0.8 (details in Methods). In each setting for each PVE value, we performed 20 simulation replicates. In each replicate, we randomly split the data into training data with 80% individuals and test data with the remaining 20% individuals. We then fitted different methods on the training data and evaluated their prediction performance on the test data. We evaluated prediction performance using either the squared correlation coefficient (R^2^) or mean squared error (MSE). We contrasted the prediction performance of all other methods with that of ISR by taking the difference of R^2^ or MSE between the other methods and ISR. Therefore, an R^2^ difference below zero or an MSE difference above zero suggests worse performance than ISR. For each result of the box plot, it consists of five numerical points: minimum (lower edge), lower quartile (25%, Q1), median (solid line in the box), upper quartile (75 %, Q3), and maximum value (upper edge). The lower quartile, median, and upper quartile form a box with compartments. An extension line is established between the upper quartile and the maximum value. This extension line is called a “whisker”. Since there are always large differences in the values, these deviating data points are listed separately in the figure (the blue points in the figure), so the whiskers in the figure can be modified to the smallest observation value and the largest observation in two levels Value, that is, the maximum observation value (max = *Q3* – 1.5 ×*IQR*) and the minimum observation value (min = *Q*1+1.5*×IQR*) is set to 1.5 IQR (interquartile range) of the distance between the quartile value.

Figure 2 shows R^2^ and MSE differences for different methods across 20 replicates in each of the four simulation settings for PVE = 0.5. Because Fig. 2 shows prediction performance difference, a large sample variance of a method in the figure only implies that the prediction performance of the method differs a lot from that of ISR, but does not imply that the method itself has a large variation in predictive performance. Supplementary Table 1 shows the means and the standard deviation of absolute R^2^ values across cross variation replicates; various methods display similar prediction variability. Supplementary Figs. 1 and 2 show the R^2^ and MSE differences for PVE = 0.2 and PVE = 0.8, respectively. The R^2^ and MSE values of the baseline method, ISR, are shown in the corresponding figure legend.

**Fig. 2.**
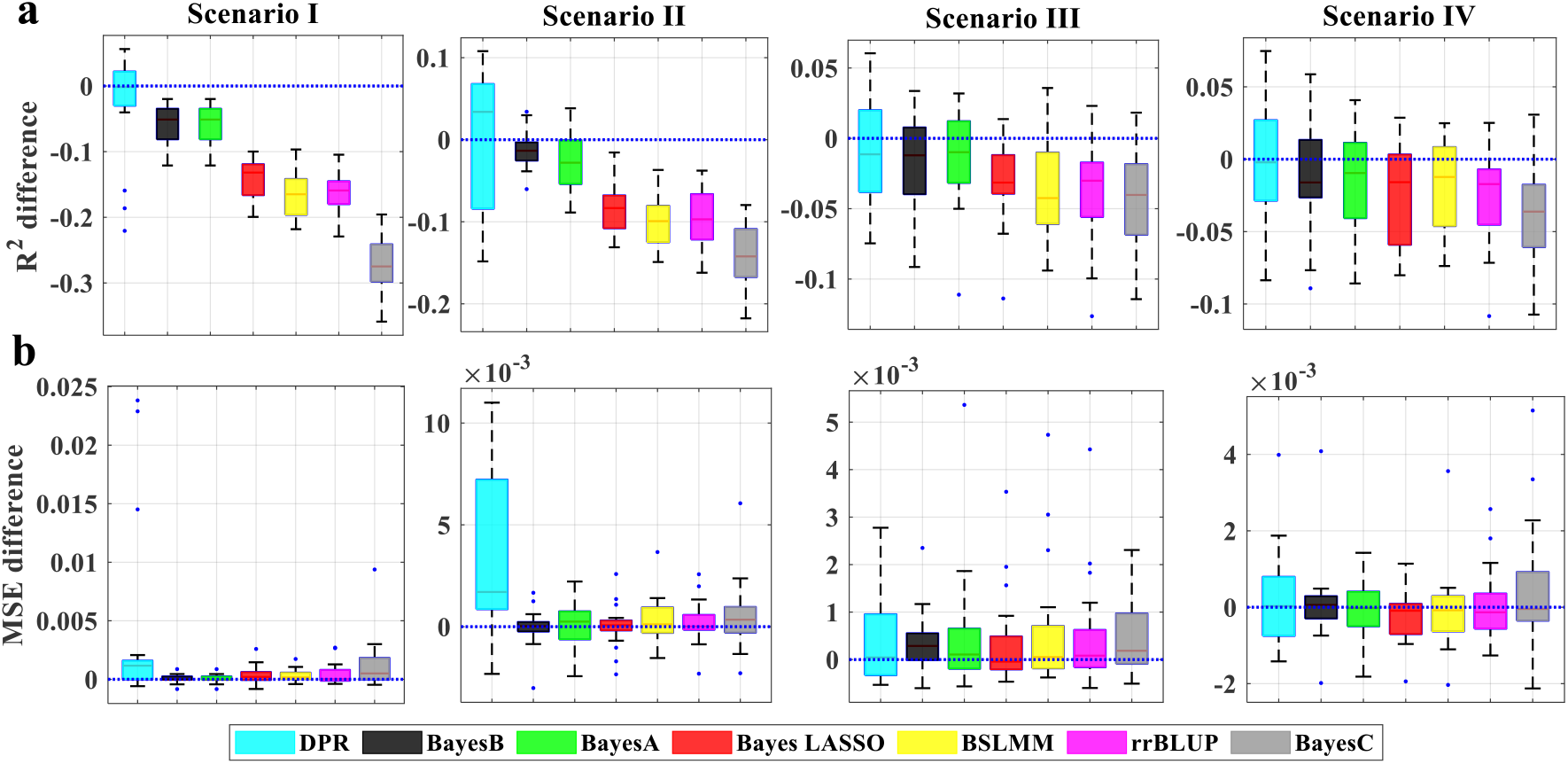
Comparison of prediction performance of seven methods with ISR in simulations when PVE = 0.5. Performance is measured by R^2^ difference (a) and MSE difference (b) with respect to ISR, where an R^2^ difference below zero (i.e., values below the blue horizontal line) or an MSE difference above zero suggests worse performance than ISR. The sample R2 and MSE differences are obtained from 20 replicates in each scenario. Methods for comparison include DPR (cyan), BayesB (black), BayesA (green), Bayes LASSO (red), BSLMM (yellow), rrBLUP (purple), and BayesC (gray). Simulation scenarios include Scenario I, Scenario II, and Scenario III, which satisfies the DPR modeling assumption; where the number of SNPs in the large effect group is 100, 150, or 500; and Scenario IV, which satisfies the BayesR modeling assumption; For each box plot, the bottom and top of the box are the first and third quartiles, while the ends of whiskers represent either the lowest datum within 1.5 interquartile range of the lower quartile or the highest datum within 1.5 interquartile range of the upper quartile. For ISR, the mean predictive R^2^ in the test set and the standard deviation for the eight settings are, respectively, 0.441 (0.019), 0.331 (0.028), 0.267 (0.016), 0.271 (0.023)

As in the previous study shown^19^, each method works the best when their modeling assumption is satisfied. In our study also shown that ISR is robust and performs well and stabilization across all twelve settings from four scenarios. For example, if we rank the methods based on their median of R2 and MSE difference (boxplot red line) performance across replicates, then when the total PVE is moderate (e.g., PVE = 0.5, Fig. 2; note that for each PVE there are a total of four simulation settings for the four scenarios), are the best or among the best (where “among the best” refers to the case when the difference between the given method and the best method is within ± 0.005 with ISR) in four simulation settings. Similarly, when the total PVE is high (e.g., PVE = 0.8, Supplementary Fig. 2), ISR is the best or among the best in four simulation settings and performance more stabilization in four simulation settings, and it is ranked as the second-best in scenario II which based on Scenario I that we appended 50 SNPs to group-three SNPs. Even when ISR is ranked as the second-best method, the difference between ISR and the best method is often small. Among the rest of the methods, BSLMM, BayesA, BayesLASSO, rrBLUP, BayesB, BayesC all work well in polygenic settings (e.g., PVE = 0.2, Supplementary Fig. 1, scenario I, scenario III, and scenario IV) but can perform poorly in sparse settings with high PVE (e.g., PVE = 0.8, Supplementary Fig. 2). The performance of DPR and BSLMM in polygenic vs. sparse settings presumably stems from their polygenic assumptions on the effect size distribution. In contrast, because of the sparse assumption on the effect size distribution, DPR has an advantage in sparse settings (e.g., PVE = 0.8, Supplementary Fig. 2; scenario III and scenario IV) but the performance of DPR is also generally worse than ISR in the challenging setting when PVE is either small or moderate, presumably because of the much simpler prior assumption employed in BVSR for the non-zero effects.

### Real data applications

To gain further insights, we compare the performance of ISR with the other methods in four real data sets to perform genomic selection in animal and plant studies.

We compare the performance of ISR with the other methods in predicting phenotypes in three GWAS data sets: (1) a cattle study^25^, where we focus on three phenotypes: milk fat percentage (MFP), MY, as well as somatic cell score (SCS); (2) a rice study^43^, where we use GL as the phenotype; (3) the Carworth Farms White (CFW) data^44^, where we focus on ten traits that include that the heritability estimates are:0.49 testweight (testes weight), 0.28 for soleus, 0.25 for plantaris, 0.10 for fastglucose (fasting glucose), 0.41 for tibial (tibia length), 0.60 for BMD (Bone-mineral density), 0.39 for TA (tibialis anterior), 0.37 for EDL (extensor digitorum longus), 0.25 for gastric (gastrocnemius), and 0.29 for sacweight (Testis weights). (4) Wheat PHS data^45^. As in simulations, for each phenotype, we performed 20 Monte Carlo cross validation data splits, except for the wheat PHS data. In each data split, we fitted methods in a training set with 80% of randomly selected individuals and evaluated method performance using R^2^ or MSE in a test set with the remaining 20% of individuals. Because the wheat PHS data set is small, we use the 10-fold cross validation method to analyze the predictive power of different methods, which is to randomly divide the sample into ten equal parts each time, and nine of them are used as training samples. The other one is used as a verification sample, and nine samples are used to estimate the parameters to predict the remaining one, and the loop 10 times in turn until all individuals are predicted. We again contrasted the performance of the other methods with that of ISR by taking the R^2^ difference or MSE difference with respect to ISR. The results are shown in Fig. 3 (R^2^ difference) and Supplementary Fig. 3 (MSE difference), with R^2^ and MSE of ISR presented in the corresponding figure legend. Supplementary Table 1 shows the means and standard deviation of absolute R^2^ values across cross variation replicates.

**Fig. 3.**
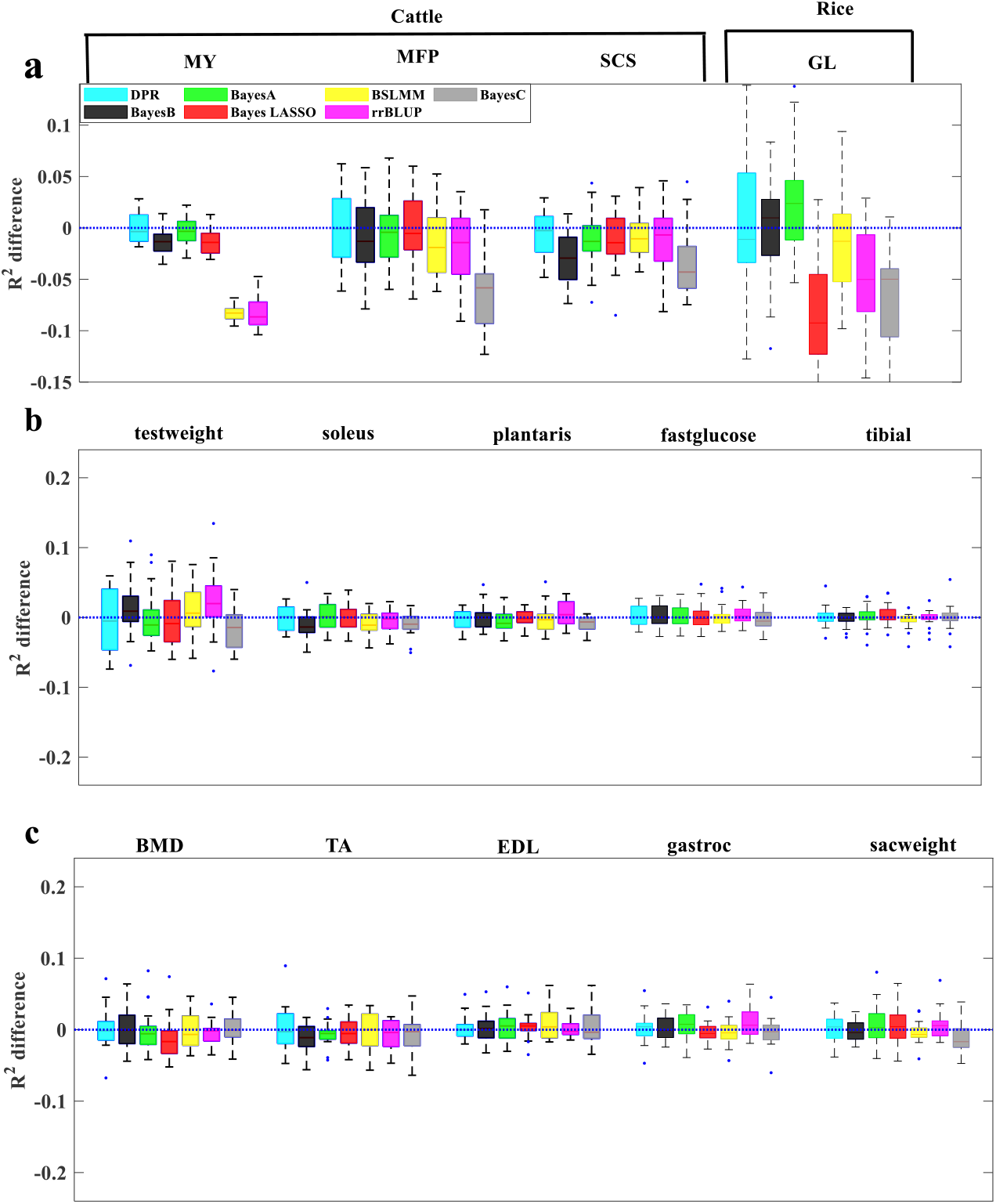
Comparison of prediction performance of seven methods with ISR for fourteen traits from three data sets. a Prediction performance for MFP, MY, and SCS in the cattle data, and for GL in the rice data; b,c Prediction performance for the ten traits in the mice data. Performance is measured by R^2^ difference with respect to ISR, where a negative value (i.e., values below the red horizontal line) indicates worse performance than ISR. Methods for comparison include DPR (cyan), BayesB (black), BayesA (green), Bayes LASSO (red), BSLMM (yellow), rrBLUP (purple), and BayesC (gray). For each box plot, the bottom and top of the box are the first and third quartiles, while the ends of whiskers represent either the lowest datum within 1.5 interquartile range of the lower quartile or the highest datum within 1.5 interquartile range of the upper quartile. The sample R^2^ differences are obtained from 20 replicates of Monte Carlo cross-validation for each trait. For ISR, the mean predictive R^2^ in the test set and the standard deviation across replicates are 0.747(0.007) for MFP, 0.618(0.03) for MY, 0.554(0.018) for SCS and 0.658(0.032) for GL, 0.078(0.024) for testweight, 0.024(0.012) for soleus, 0.017(0.011) for plantaris, 0.018(0.009) for fastglucose, 0.008(0.01) for tibial, 0.057(0.008) for BMD, 0.028(0.016) for TA, 0.022(0.005) for EDL, 0.02(0.013) for gastric, and 0.028(0.01) for sacweight. The heritability estimates are 0.912 for MFP, 0.810 for MY, 0.801 for SCS, and 0.976 for GL, 0.49 for testweight, 0.28 for soleus, 0.25 for plantaris, 0.10 for fastglucose, 0.27 for tibial, 0.60 for BMD, 0.39 for TA, 0.37 for EDL, 0.25 for gastric, and 0.29 for sacweight.

Overall, consistent with simulations, ISR shows robust performance across all traits and is ranked either as the best or the second-best method or equivalent. In the cattle data (Fig. 2a), for SCS and MY, both ISR and DPR perform the best. For MFP, ISR and DPR perform equivalent, followed BayesA, BayesB, BayesLASSO, BSLMM, rrBLUP, and BayesC. while BSLMM and rrBLUP do not perform well for MY in the cattle data, but their performance improves for MFP and SCS, consistent with scenario III and scenario IV (simulation hypothesis is constant). The relative performance of ISR, DPR BayesA, BayesB in the cattle data is compatible with the distinct genetic architectures that underlie the three complex traits^25,46^. While MFP and MY are affected by a few large or moderate effect SNPs and many small effect SNPs, SCS is a highly polygenic trait influenced by many SNPs with small effects. BayesC performs poorly for these three traits in the cattle data. In the rice data (Fig. 2a), BayesA performs the best, followed by ISR, PDR, BayesB, BSLMM, rrBLUP, BayesC, BayesLASSO, suggesting that a few SNPs influence GL with large effects^43^. In the CFW data (Fig. 2b, c), ISR performs the best or among the best for testweight, soleus, plantaris, BMD, and TA. Its performance is comparable to BayesB and rrBLUP for plantaris, and follows right behind DPR. Its also performance is comparable to DPR, BayesA, BayesB, and rrBLUP for EDL, gastric, and sacweight, and follows right behind BSLMM. However, it can be seen from the MSE difference that compared with ISR, the performance is poor, and its value is above 0, indicating that the predictive power of this method is quite different, although there may be several times in the 20 cross-validations A large predictive power can be obtained. Both the CFW phenotype was low PVE^44^.

Because the wheat PHS is a family-based study that PHS resistance showed varied effects under different environments^45^. The wheat PHS resistance traits are rarely used in genome selection and evaluated (prediction) in current research. There are differences in the predictive power of different methods between different. To eliminate the environmental difference between indifference years, we have given the four-year BLUP estimate for calculation. The best performance is ISR, followed by BayesA, BayesB, BayesLASSO, rrBLUP, BSLMM, BayesC, and DPR (Supplementary Fig. 4). In each year’s data, both are ISR performs best, and followed BayesA, BayesB, BayesLASSO, and rrBLUP, BSLMM, BayesC, and DPR.

### Overview

Based on the Simulations and Real data applications (did not use the wheat PHS data) results from the averaged prediction of R^2^, we use the TOPSIS and cluster methods to ranked all methods that all-around performance (Fig.4a,b, Supplementary Table 2). Both the TOPSIS and cluster showed the same result that ISR(0.63) is perform best, and followed by DPR(0.63), BayesA(0.59), BayesLASSO(0.57), rrBLUP(0.54), BayesB(0.48), BSLMM(0.36) and BayesC(0.22). If we included the wheat dataset perform the TOPSIS and cluster analysis that also showed the ISR(0.66) is perform best, and followed by BayesB(0.57), BayesA(0.54), DPR(0.53), BayesLASSO(0.46), rrBLUP(0.46), BSLMM(0.35) and BayesC(0.19)(Supplementary Fig. 6, Supplementary Table 2,3). Finally, we list the eight methods’ computational time for the three traits only in a large dataset, the maize dataset (Supplementary Table 4). And we excluded the BayesC and added a new BayesR method. Here, we also compare predictive ability, but not described here again, as shown in the other dataset prediction results. For sampling-based methods (DPR, BayesR, BayesA, BayesB, BayesLASSO, and BSLMM), we measure the computational time based on a fixed 10,000 iterations. However, due to the different convergence properties of different algorithms, a fixed number of iterations in different methods may correspond to different mixing performance^19,20^. In contrast, rrBLUP is the faster method, while DPR, BayesR, and BSLMM are as same as computationally efficient. ISR is computationally as efficient as the other three BayesA, BayesB, and BayesLASSO for YWK, but costest time for GDD and SSK traits.

**Fig. 4.**
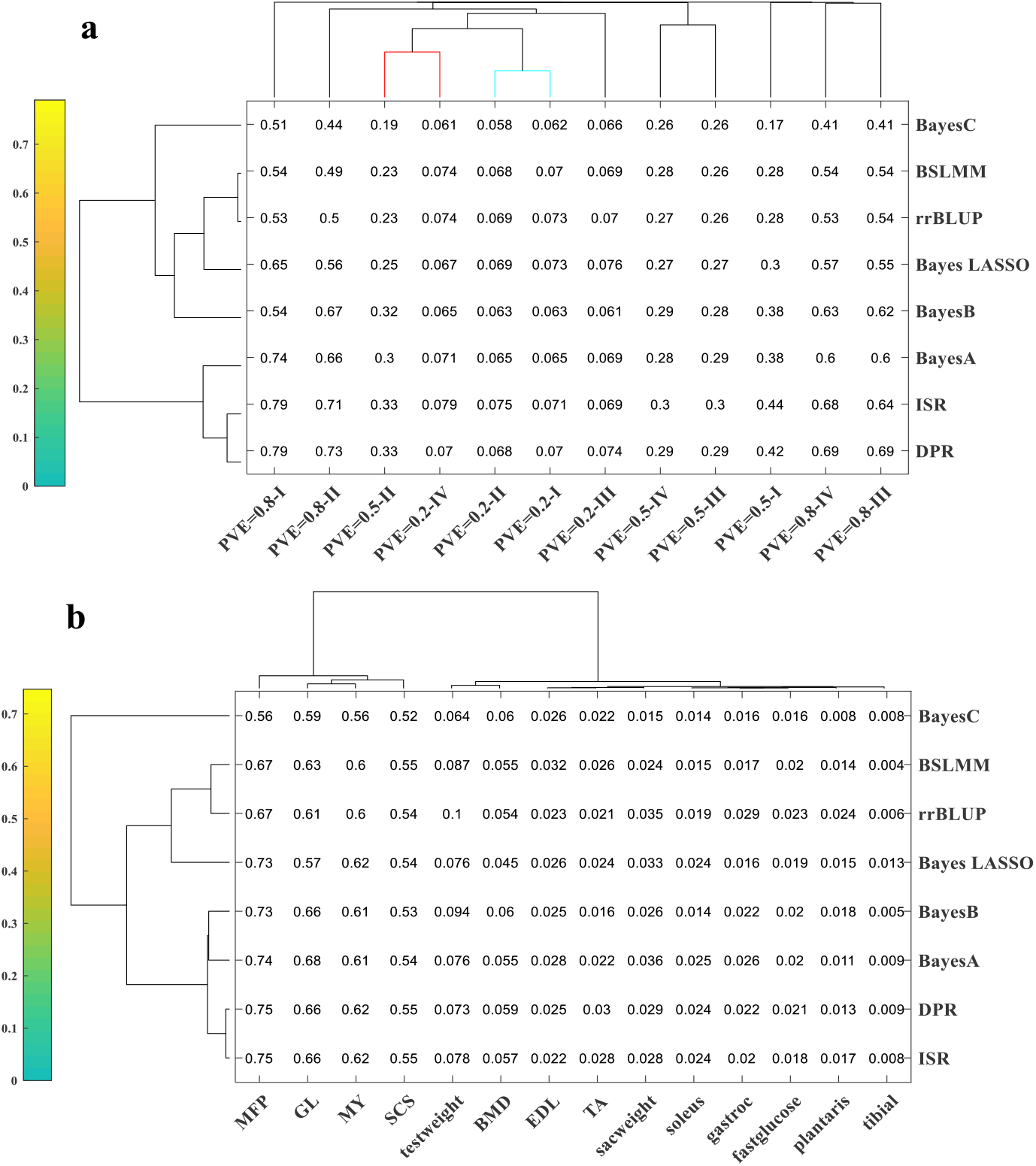
The clustering result with heatmap. Based on the Simulations and Real data applications (did not include the wheat PHS data) results in the averaged prediction of R^2^

## Discussion

We have presented a novel statistical method, ISR, for the polygenic prediction of complex traits. ISR is a flexible model for the different effect size from the normal distribution (Fig.1), which can be split into three group effects: no effect, weaker effect, and stronger effect and developed for modeling polygenic traits in genetic association studies. By flexibly modeling the difference effect size, ISR can adapt to the polygenic architecture underlying many complex features and enjoys robust performance across a range of phenotypes. With simulations and applications to five species real data sets, we have illustrated the benefits of ISR. We have focused on one application of ISR, which genetic prediction of phenotypes. As the other polygenic methods^28,40,47^, ISR can also be applied to models of traits controlled by multiple genes. For example, ISR can be used to estimate the proportion of variance in phenotypes explained by each of SNPs^42^, a quantity that is commonly referred to as SNP heritability^28,33^. Because ISR assumes a flexible effect size distribution that is adaptive to the genetic architecture underlying a given trait, it also can provide an accurate estimation of SNP heritability^42^. As another example, ISR also can be applied to association mapping (GWAS)^42^(Supplementary Fig.9,10,11, and Supplementary Table 4).

Previous studies have shown that the ISR method has a strong power to identify variant loci. It performs better than current statistical analysis methods, so we use it to perform genome-wide prediction^42^. Here, we have restricted ourselves to applying ISR to continuous phenotypes. For case-control studies (such as maize traits SSK and YWK), we could follow previous approaches of treating binary phenotypes as continuous traits and apply ISR directly^28,29,40^. In the present study, as shown in Fig.4, the cluster analysis of the predictive power of different models of simulated and real phenotypes (where the distance between variables (rows and columns are the targets) and the distance between classes are respectively used by Mahalanobis distance and the sum of squares of deviations) and found that, just as the four methods with consistent simulation results, DPR, ISR, BayesA, and BayesB performed the best, in the four different simulations at three different heritability rates, the predictive power was significant, especially at high heritability rates. It is higher than the other four methods (ANOVA, p=4.06e-07, Supplementary Fig.7), but there is no significant difference between these four methods (ANOVA, p=0.1403 Supplementary Fig.7). Under the moderate heritability, the average predictive power of ISR is the highest. However, except that it is significantly higher than BayesC (ANOVA, p=0.043, Supplementary Fig.7) and the remaining methods have no statistically significant differences; as the same, BayesA has the highest average predictive power at low heritability, and the same except that it is significantly higher than (ANOVA, p=0.0141, Supplementary Fig.7) The difference between the outer BayesC and the remaining methods is not significant (ANOVA, p=0.0858, Supplementary Fig.7), which is consistent with the result analysis (Fig.1, Supplementary Fig.1,2). In addition, the classification given by the cluster analysis between the columns is also very reasonable (Fig.4a, the different colors of the cluster tree).

The true phenotype analysis is also showed the same with simulation, dividing different predictive powers into four categories from low to high (Figs.4b, different colors of cluster trees). According to previous studies, the heritability of the three traits of the for cattle species is 0.94, 0.95, and 0.88 ^25^; the grain length of rice is 0.976^43^; the germination rate of wheat is 0.83^45^. The difference between field and greenhouse experiments is 0.92 and 0.62. The proportion of variance in phenotypes explained (PVE) of the ten traits of the remaining mice is0.49 for testweight, 0.28 for soleus, 0.25 for plantaris, 0.10 for fastglucose, 0.27 for tibial, 0.60 for BMD, 0.39 for TA, 0.37 for EDL, 0.25 for gastric, and 0.29 for sacweight^44^. It was found that all phenotypes can be grouped into four categories according to their PVE rate. For cattle, ISR and DPR have the highest average predictive ability, but there is no significant difference among the BayesA and BayesB methods (ANOVA, p=0.7314). This result is consistent with simulation Fig.2, which also shows that the differences between MSE value can explain the difference that the accuracy of difference prediction methods^19,28,31,40^; In contrast, BayesA, BayesB, and ISR have the highest predictive power in wheat PHS-2012 dataset, and they are significantly higher than other methods (ANOVA, p =0.0133); and the highest predictive ability of the remaining wheat PHS is ISR, But has no difference compared with the rest of the method (ANOVA, p=0.976, Supplementary Fig.8). Here, we can find out that the estimated value of BLUP in four years which has the highest predictive abillity is ISR, where is similar to Moore et al.’s research used the marker-assisted selection (0.40~0.59)^48^; While with the low of PVE (heritability) of CFW dataset, there are no difference in predictive ability between methods (ANOVA, p=0.998, Supplementary Fig.8), but the ISR and DPR always has the highest average pretictive ability (Supplementary Table 1).

In a words, the performance of all methods in simulating and real phenotype-wide prediction is consistent (performance under different heritability (PVE)). Therefore, here we use the TOPSIS^49^ comprehensive evaluation method, which combining the averages of predictive ability of the simulation and real phenotypes as variables, and the goal is to rank all methods comprehensively. Where the result show that ISR(0.63) is perform best, and followed by DPR(0.63), BayesA(0.59), BayesLASSO(0.57), rrBLUP(0.54), BayesB(0.48), BSLMM(0.36) and BayesC(0.22). While considered the wheat PHS dataset was small and affected by more the environment with different years (Supplementary Fig. 5, Supplementary Table 2,3).

Of course, this study only analyzes the traits related to animals and plants and does not analyze human diseases related to features (conditional restrictions). Human studies are based on tens of thousands of individuals and millions of genetic markers, just like Zeng et al.’s simulation and disease real phenotype research showed that the result currently DPR and BayesR were relatively best prediction methods^19^. In addition, since the control of human diseases is mainly controlled by many genes and many minor genes (many genetic markers with small effects) ^8,50,51^, they also can reasonably estimate the effective SNP PVE (narrow-sense heritability)^51,52^. DPR, which is consistent with the results of the simulation study by Zeng et al^19^, and was indeed superior to other methods (Fig.2). However, the complex posterior distributions and computational complexity of traditional multiple integrals limited Bayesian methods^20^. The problem was solved after the MCMC method and the Gibbs algorithm were introduced to Bayesian statistics. However, in condition M(SNPs)≫N (samples), which MCMC and Gibbs algorithm iterations is hard to reach the convergence of the posterior means, which limits the practical application of Bayesian methods^9,28,40,53,54^.

The ISR method is not without its defects. In addition to the calculated efficiency (Supplementary Table 4), if the trait is controlled by many genes and minor genes (all SNPs genetic markers have smaller effects), then there will be cases where the predictive ability is low (Fig.1, Supplementary Fig.1,2,5). For example, the predictive power was low when simulating 500 SNPs (under low to medium heritability). However, our ISR model can fit the epistasis effect, where if the interaction between genes is considered, its predictive ability will be improved^55-57^. Although the simulation and real performance results show that ISR is superior to other models(Supplementary Fig. 6, Supplementary Table 2,3), there is still a lot of room for improvement in this polygenic prediction model. For example, the algorithm’s improvement, combined with the optimization of the model objective function, can make the ISR perform better. The complexity of the calculation time also needs to be optimized.

## Methods

### Overview of ISR

We provide a brief overview of ISR here. Detailed methods and algorithms are provided^42^. To model the relationship between phenotypes and genotypes, we consider the following multiple regression model:

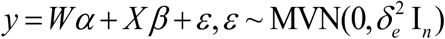

where *y* is an *n*-vector of phenotypes measured on *n* individuals; W=(w_1_, w_2_… w_c_) is an *n* by *c* matrix of covariates(fixed effects) including a column of ones for the intercept term; *α* is a *c*-vector of coefficients; X is an *n* by *p* matrix of genotypes; β is the corresponding *p*-vector of effect sizes; ε is an *n*-vector of residual errors where each element is assumed to be independently and identically distributed from a normal distribution with a variance 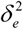; In is an *n* by *n* identity matrix and MVN denotes multivariate normal distribution.

We used the proposed iterative screening regression model—effect size estimates obtained by the least-square method (LSM) and F-test P values for each SNP. The SNP with the most significant association is then added to the model as a cofactor for the next step. Combined the proposed iterative screening regression process, which makes it useful when*p≫n* (when the number of SNPs is much greater than the number of individuals). We also proposed a new model selection criteria (RIC Fig.1) to select the most appropriate model^42^.

### Simulations

We used genotypes from an existing cattle GWAS data set with 5024 individuals and 42,551 SNPs and simulated phenotypes. To cover a range of possible genetic architectures, we consider four different simulation scenarios to cover a range of possible genetic architectures:

Scenario I, where we randomly selected 100 SNPs, are causal and SNPs in different effect-size groups have different effects. Specifically, we randomly selected 10 group-one SNPs, 40 group-two SNPs, 50 group-three SNPs, and set the remaining SNPs to have zero effects. We simulated SNP effect sizes all from a standard normal distribution but scaled their effects in each group separately so that the proportion of genetic variance explained by the four groups are 0.15, 0.25, and 0.60, respectively. We set the total proportion of phenotypic variance (PVE; i.e., SNP heritability) to be either 0.2, 0.5, or 0.8, representing low, moderate, and high heritability, respectively. This simulation scenario consists of one simulation setting for each PVE.

Scenario II based on Scenario I that we appended 50 SNPs to group-three SNPs, the remained simulation conditions were the same. These causal SNPs come from three effect-size groups. Here, the proportion of PVE by the three groups are 0.15, 0.25, and 0.6, respectively. Again, we set the total PVE to be either 0.2, 0.5, or 0.8. This simulation scenario consists of one simulation setting for each PVE.

Scenario III is similar to Scenario I where we randomly selected 500 SNPs are causal and SNPs in different effect-size groups have different effects. Specifically, we randomly selected 50 group-one SNPs, 150 group-two SNPs, 300 group-three SNPs, and set the remaining SNPs to have zero effects. We simulated SNP effect sizes all from a standard normal distribution but scaled their effects in each group separately so that the proportion of genetic variance explained by the four groups are 0.15, 0.25, and 0.60, respectively. We set the total proportion of phenotypic variance (PVE; i.e., SNP heritability) to be either 0.2, 0.5, or 0.8, representing low, moderate, and high heritability, respectively. This simulation scenario consists of one simulation setting for each PVE.

Scenario IV satisfies the BayesR modeling assumption, where we randomly selected 500 SNPs are causal and SNPs come from three effect-size groups. Specifically, we randomly selected 50 group-one SNPs, 150 group-two SNPs, 300 group-three SNPs, and set the remaining SNPs to have zero effects. The simulated effect size follows a normal distribution with a mean value of 0 and a variance of 10^-2^, 10^-3^, and 10^-4^, respectively^40^. Here, the proportion of PVE by the three groups are 0.15, 0.25, and 0.6, respectively. Again, we set the total PVE to be either 0.2, 0.5, or 0.8. This simulation scenario consists of one simulation setting for each PVE.

To test the power of ISR method, Scenario I to Scenario III were more satisfies the ISR model, and Scenario IV satisfies the BayesR modeling assumption. Both the scenarios were as same as the real data perform. In each setting, we performed 20 simulation replicates. In each replicate, we randomly split the data into training data with 80% individuals and test data with the remaining 20% individuals. We then fitted different methods on the training data and evaluated their prediction performance on the test data.

### Cattle data

The cattle data^25^ consists of 5024 samples and 42,551 SNPs after removing SNPs that have a HWE p-value < 10^-4^, a genotype call rate <95%, or an MAF < 0.01. For the remaining SNPs, we imputed missing genotypes with the estimated mean genotype of that SNP. We analyzed three traits: MFP, MY, and SCS. All phenotypes were quantile normalized to a standard normal distribution before analysis.

### Rice data

The maize data^43^ which after processing the data, including filtering for missing genotype data which no measure the traits, and minor allele frequencies(MAF <0.05), the data were composed of m = 464,831 SNPs and n = 1,132 individuals. For the remaining SNPs, we also imputed missing genotypes with the estimated mean genotype of that SNP. We only used the grain length (GL) as the phenotype in genomic selection.

### CFW data

Outbred CFW^44^ (Carworth Farms White) mice population that including a set of 92,734 single-nucleotide polymorphism markers which were genotyped, 1,161 individuals. We analyzed ten traits: testweight, soleus, plantaris, fastglucose, tibial, BMD, TA, EDL, gastric, and sacweight. The heritability estimates are 0.49 for testweight (testes weight), 0.28 for soleus, 0.25 for plantaris, 0.10 for fastglucose (fasting glucose), 0.41 for tibial (tibia length), 0.60 for BMD (Bone-mineral density), 0.39 for TA (tibialis anterior), 0.37 for EDL (extensor digitorum longus), 0.25 for gastric (gastrocnemius), and 0.29 for sacweight (Testis weights)^44^.

### Wheat PHS data

A set of 185 winter wheat accessions^45^, and included 27521 SNPs. The GWAS panel was evaluated for PHS in the greenhouse experiments of fall (August-December) 2011, spring (January-May) and fall 2012, and spring 2013. All experiments were conducted in a randomized complete block design with two replications of five plants. The GWAS panel was also planted for PHS resistance evaluation in the Kansas State University Rocky Ford Wheat Research Farm, Manhattan, KS and the Agricultural Research Center-Hays, Hays, KS, respectively, in the summers of 2013 and 2014. PHS values of four years were used for BLUP estimation to obtain BLUP values for prediction analysis. The broad-sense heritability across all experiments was high (0.83), with 0.62 in the greenhouse experiments and 0.92 in the field experiments^45^.

### Maize data

As described^20,58^ that the maize data consisted of 2279 inbred accessions and three traits, including two case/control traits: yellow or white kernels YWK) and sweet or starchy kernels (SSK), and one quantitative trait: growing degree days (GDD). A total of 681,257 SNPs across all maize lines were obtained with genotyping by sequencing (GBS). After removing samples missing is > 20%, SNPs with either MAF < 0.01, 2279 individuals and 195,038 SNPs for GDD; 314 controls, 1281 cases, and 185,493 SNPs for YWK; 2490 controls, 141 cases, and 183,225 SNPs for SSK; remained in this study. We imputation the missing genotype data with Beagle5.1 (https://faculty.washington.edu/browning/beagle/beagle.html)^59,60^. And we perform the GWAS use the ISR model only (Supplementary Fig 9,10,11, Supplementary Table 5). The PVE estimates are 0.88 for GDD, 0.63 for SSK, 0.97 for YWK.

### Other methods

We compared the performance of ISR mainly with seven existing methods: (1) DPR^19^; (2) BSLMM (implemented in the GEMMA software (version 0.95alpha))^28^; (3) BayesA; (4) BayesB; (5) BayesC; (6) Bayes LASSO; (7) rrBLUP^35^, (8) BayesR^40^. Among them (3)-(6) the method of receiving in BGLR R package. We used default settings to fit all these methods. To measure prediction performance, we carried out 20 Monte Carlo cross-validation data splits as in simulations. In each data split, we fitted methods in a training set with 80% of randomly selected individuals and evaluated method performance using R^2^ in the test set with the remaining 20% of individuals. Because the wheat data set is small, we use the 10-fold cross-validation method to analyze the predictive power of different methods, which is to divide the sample into ten equal parts each time randomly, and nine of them are used as training samples. The other one is used as a verification sample, and nine samples are used to estimate the parameters to predict the remaining one, and the loop 10 times in turn until all individuals are predicted.

### TOPSIS method

TOPSIS (Technique for Order Preference by Similarity to Ideal Solution), the technique of approximating the ideal solution, is a multi-criteria decision analysis method. The basic idea of this method is to define the ideal solution and the negative ideal solution of the decision-making problem. After the ideal solution and the negative ideal solution are determined, the distance between the evaluation object and the optimal solution and the worst solution is calculated respectively, so as to obtain and the optimal solution through calculation. If a certain evaluation object is infinitely close to the ideal solution and at the same time far away from the negative ideal solution, then this solution is the optimal solution.

How to calculate the distance is very important. The TOPSIS method uses the Euclidean distance function to calculate the distance between the evaluation object and the ideal solution and the negative ideal solution. The Euclidean distance describes the true distance between two points in the p-dimensional space. Here, suppose there are two points in space *A* = (*a*_1_, *a*_2_,…, *a_n_*) and *B* = (*b*_1_, *b*_2_,…, *b_n_*), then, The Euclidean distance calculation formula is as follows:

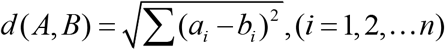

Suppose the sample material is a multi-attribute decision-making matrix with *n* evaluation objects and *m* evaluation indicators The TOPSIS process is carried out as follows:

Step 1: Convergence processing for each index of the sample material. As the evaluation process requires the same trend of indicators, that is, either the higher the better, or the lower the better. Therefore, the original data needs to be converted, that is, the conversion of low-quality indicators to high-quality indicators or the conversion of high-quality indicators to low-quality indicators.

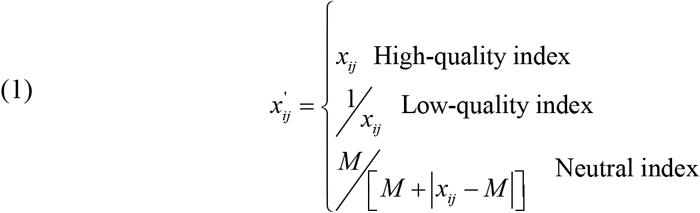
Construct a normalized decision matrix. In the target decision-making, the different dimensions of the evaluation index will have a great impact on the evaluation result. The range of changes of each index is different, and there is no unified measurement standard. Therefore, the decision matrix needs to be normalized.

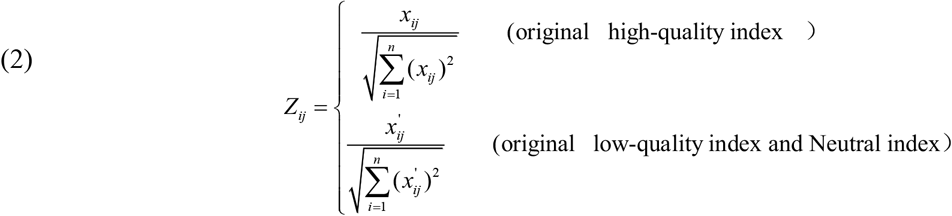
Step 3: Find the best plan and the worst plan:

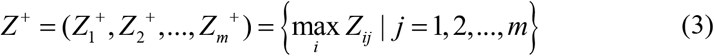

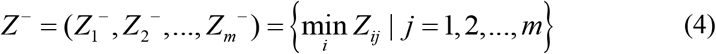
Step 4: Calculate the Euclidean distance between each evaluation object and the ideal solution and the negative ideal solution.

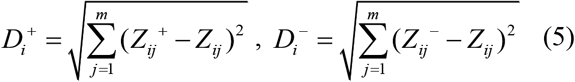 In the formula, 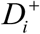 and 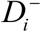 respectively represent the distance between the *i*-th evaluation object and the ideal solution and the negative ideal solution; represent the *j*-th index data of the *i*-th material in the normalized matrix.
Step 5: Calculate the closeness of *C_i_* each target solution to the optimal solution to reflect the quality of the target solution.

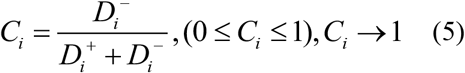
Step 6: Sort by size *C_i_* and give the evaluation result. The larger the value of *C_i_*, the better the overall benefit and the better the plan.

### Cluster method

Here, we used hierarchical clustering to evaluate the different methods perform and use the heat map with dendrograms to show the result. Algorithm for computing the distance between clusters that we use the ward method and the distance metric was calculated by Mahalanobis distance, as follows:

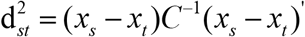

where *C* is the covariance matrix. Mahalanobis distance is widely used in cluster analysis and classification techniques. It is closely related to Hotelling’s T-square distribution used for multivariate statistical testing and Fisher’s Linear Discriminant Analysis that is used for supervised classification^61^.

## Code availability

Our method is implemented in the ISR software included TOPSIS and cluster methods, and all script methods analysis in this study can freely available at https://github.com/czheluo/PPISR and https://github.com/czheluo/ISR.

## Data availability

No data were generated in the present study. The genotype and phenotype data from the Cattle from^25^ and Cattle: https://www.g3journal.org/content/5/4/615. supplemental; and Maize: https://datacommons.cyverse.org/browse/iplant/home/shared/panzea. And rice data studies are available http://www.ricediversity.org/data/. The outbred CFW mice of genotype and phenotype data are publicly available at https://github.com/pcarbo/cfw, and the genotype was as same as the Parker, C.C et.^42,44^ and the wheat PHS data set provided by Prof. Guihua Bai at the Kansas State University.

## Author contributions

Shiliang Gu and Meng Luo conceived the study and supervised statistical aspects and developed the algorithm of this work, and developed the software. Meng Luo designed the experiment and performed the simulations and data analyses. Meng Luo wrote the manuscript.

## Additional information

Supplementary Information accompanies this paper.

## References

1. Makowsky, R. et al. Beyond Missing Heritability: Prediction of Complex Traits. PLOS Genetics 7, e1002051 (2011).

2. Millet, E.J., Kruijer, W., Coupel-Ledru, A., Prado, S.A. & Tardieu, F. Genomic prediction of maize yield across European environmental conditions. Nature Genetics 51(2019).

3. Wray, N.R. et al. Pitfalls of predicting complex traits from SNPs. Nature Reviews Genetics 14, 507 (2013).

4. Hayes, B.J., Pryce, J., Chamberlain, A.J., Bowman, P.J. & Goddard, M.E. Genetic Architecture of Complex Traits and Accuracy of Genomic Prediction: Coat Colour, Milk-Fat Percentage, and Type in Holstein Cattle as Contrasting Model Traits. PLOS Genetics 6, e1001139 (2010).

5. Georges, M., Charlier, C. & Hayes, B. Harnessing genomic information for livestock improvement. Nature Reviews Genetics 20, 135–156 (2019).

6. Desta, Z.A. & Ortiz, R. Genomic selection: genome-wide prediction in plant improvement. Trends in Plant Science 19, 592–601 (2014).

7. Khera, A.V. et al. Polygenic Prediction of Weight and Obesity Trajectories from Birth to Adulthood. Cell 177, 587–596. e9 (2019).

8. Chatterjee, N., Shi, J. & García-Closas, M. Developing and evaluating polygenic risk prediction models for stratified disease prevention. Nature Reviews Genetics 17, 392–406 (2016).

9. Ge, T., Chen, C.-Y., Ni, Y., Feng, Y.-C.A. & Smoller, J.W. Polygenic prediction via Bayesian regression and continuous shrinkage priors. Nature Communications 10, 1776 (2019).

10. Khera, A.V. et al. Genome-wide polygenic scores for common diseases identify individuals with risk equivalent to monogenic mutations. Nature Genetics 50, 1219–1224 (2018).

11. Maier, R.M. et al. Improving genetic prediction by leveraging genetic correlations among human diseases and traits. Nature Communications 9, 989 (2018).

12. Pasaniuc, B. & Price, A.L. Dissecting the genetics of complex traits using summary association statistics. Nature Reviews Genetics 18, 117–127 (2017).

13. Chatterjee, N., Shi, J. & García-Closas, M. Developing and evaluating polygenic risk prediction models for stratified disease prevention. Nature Reviews Genetics 17, 392 (2016).

14. Wiggans, G.R., Cole, J.B., Hubbard, S.M. & Sonstegard, T.S. Genomic Selection in Dairy Cattle: The USDA Experience. Annual Review of Animal Biosciences 5, 309–327 (2017).

15. Crossa, J. et al. Genomic Selection in Plant Breeding: Methods, Models, and Perspectives. Trends in Plant Science 22, 961–975.

16. Lozada, D.N., Mason, R.E., Sarinelli, J.M. & Brown-Guedira, G. Accuracy of genomic selection for grain yield and agronomic traits in soft red winter wheat. BMC Genetics 20, 82 (2019).

17. Ali, M., Zhang, Y., Rasheed, A., Wang, J. & Zhang, L. Genomic Prediction for Grain Yield and Yield-Related Traits in Chinese Winter Wheat. International Journal of Molecular Sciences 21(2020).

18. Gamazon, E.R. A gene-based association method for mapping traits using reference transcriptome data. Nat. Genet. 47(2015).

19. Zeng, P. & Zhou, X. Non-parametric genetic prediction of complex traits with latent Dirichlet process regression models. Nature Communications 8, 456 (2017).

20. Yin, L. et al. KAML: improving genomic prediction accuracy of complex traits using machine learning determined parameters. Genome Biology 21, 146 (2020).

21. Lango Allen, H. et al. Hundreds of variants clustered in genomic loci and biological pathways affect human height. Nature 467, 832 (2010).

22. Locke, A.E. et al. Genetic studies of body mass index yield new insights for obesity biology. Nature 518, 197–206 (2015).

23. Huang, X. et al. Genome-wide association studies of 14 agronomic traits in rice landraces. Nature Genetics 42, 961–967 (2010).

24. Fernandes Júnior, G.A. et al. Genomic prediction of breeding values for carcass traits in Nellore cattle. Genetics Selection Evolution 48, 7 (2016).

25. Zhang, Z. et al. Accuracy of Whole-Genome Prediction Using a Genetic Architecture-Enhanced Variance-Covariance Matrix. G3: Genes/Genomes/Genetics 5, 615 (2015).

26. Meuwissen, T., Hayes, B. & Goddard, M. Accelerating Improvement of Livestock with Genomic Selection. Annual Review of Animal Biosciences 1, 221–237 (2013).

27. Klasen, J.R. et al. A multi-marker association method for genome-wide association studies without the need for population structure correction. 7, 13299 (2016).

28. Zhou, X., Carbonetto, P. & Stephens, M. Polygenic Modeling with Bayesian Sparse Linear Mixed Models. PLOS Genetics 9, e1003264 (2013).

29. Speed, D. & Balding, D.J. MultiBLUP: improved SNP-based prediction for complex traits. Genome Research 24, 1550–1557 (2014).

30. Chatterjee, N. et al. Projecting the performance of risk prediction based on polygenic analyses of genome-wide association studies. Nature Genetics 45, 400 (2013).

31. Weissbrod, O., Geiger, D. & Rosset, S. Multikernel linear mixed models for complex phenotype prediction. Genome Research 26, 969–979 (2016).

32. Shah, S. et al. Improving Phenotypic Prediction by Combining Genetic and Epigenetic Associations. The American Journal of Human Genetics 97, 75–85.

33. Yang, J. et al. Common SNPs explain a large proportion of the heritability for human height. Nature Genetics 42, 565–569 (2010).

34. VanRaden, P.M. Efficient Methods to Compute Genomic Predictions. Journal of Dairy Science 91, 4414–4423 (2008).

35. Endelman, J.B. Ridge Regression and Other Kernels for Genomic Selection with R Package rrBLUP. The Plant Genome 4, 250–255 (2011).

36. Habier, D., Fernando, R.L., Kizilkaya, K. & Garrick, D.J. Extension of the bayesian alphabet for genomic selection. BMC Bioinformatics 12, 186 (2011).

37. Meuwissen, T.H., Hayes, B.J. & Goddard, M.E. Prediction of total genetic value using genome-wide dense marker maps. Genetics 157, 1819–1829 (2001).

38. Park, T. & Casella, G. The Bayesian Lasso. Journal of the American Statistical Association 103, 681–686 (2008).

39. Yi, N. & Xu, S. Bayesian LASSO for Quantitative Trait Loci Mapping. Genetics 179, 1045 (2008).

40. Moser, G. et al. Simultaneous Discovery, Estimation and Prediction Analysis of Complex Traits Using a Bayesian Mixture Model. PLOS Genetics 11, e1004969 (2015).

41. Cheng, W., Ramachandran, S. & Crawford, L. Estimation of non-null SNP effect size distributions enables the detection of enriched genes underlying complex traits. PLOS Genetics 16, e1008855 (2020).

42. Luo, M. & Gu, S. A new approach of dissecting genetic effects for complex traits. bioRxiv, 2020.10.16.336180 (2020).

43. McCouch, S.R. et al. Open access resources for genome-wide association mapping in rice. Nature Communications 7, 10532 (2016).

44. Parker, C.C. et al. Genome-wide association study of behavioral, physiological and gene expression traits in outbred CFW mice. Nature Genetics 48, 919 (2016).

45. Lin, M. et al. Genome-wide association analysis on pre-harvest sprouting resistance and grain color in U.S. winter wheat. BMC Genomics 17, 794 (2016).

46. Hu, Z.-L., Park, C.A., Wu, X.-L. & Reecy, J.M. Animal QTLdb: an improved database tool for livestock animal QTL/association data dissemination in the post-genome era. Nucleic Acids Research 41, D871–D879 (2013).

47. Carbonetto, P. & Stephens, M. Stephens M: Scalable variational inference for Bayesian variable selection in regression, and its accuracy in genetic association studies. Bayesian Analysis. Bayesian Analysis 7, 73–107 (2013).

48. Moore, J.K. et al. Improving Genomic Prediction for Pre-Harvest Sprouting Tolerance in Wheat by Weighting Large-Effect Quantitative Trait Loci. Crop Science 57, 1315–1324 (2017).

49. Yoon, K. A Reconciliation Among Discrete Compromise Solutions. Journal of the Operational Research Society 38, 277–286 (1987).

50. Vilhjálmsson, Bjarni J. et al. Modeling Linkage Disequilibrium Increases Accuracy of Polygenic Risk Scores. The American Journal of Human Genetics 97, 576–592 (2015).

51. Yang, J., Zeng, J., Goddard, M.E., Wray, N.R. & Visscher, P.M. Concepts, estimation and interpretation of SNP-based heritability. Nature Genetics 49, 1304 (2017).

52. Speed, D. et al. Reevaluation of SNP heritability in complex human traits. Nature Genetics 49, 986 (2017).

53. Zhu, X. & Stephens, M. Bayesian large-scale multiple regression with summary statistics from genome-wide association studies. Annals of Applied Statistics 11, 1561–1592 (2016).

54. Muller, P. & Mitra, R. Bayesian Nonparametric Inference - Why and How. Bayesian Anal. 8, 269–302 (2013).

55. Martini, J.W.R. et al. Genomic prediction with epistasis models: on the marker-coding-dependent performance of the extended GBLUP and properties of the categorical epistasis model (CE). BMC Bioinformatics 18, 3 (2017).

56. Akdemir, D., Jannink, J.L. & Isidrosánchez, J. Locally epistatic models for genome-wide prediction and association by importance sampling. Genetics Selection Evolution 49, 74 (2017).

57. Forneris, N.S., Vitezica, Z.G., Legarra, A. & Pérez-Enciso, M. Influence of epistasis on response to genomic selection using complete sequence data. Genetics Selection Evolution 49, 66 (2017).

58. Romay, M.C. et al. Comprehensive genotyping of the USA national maize inbred seed bank. Genome Biology 14, R55 (2013).

59. Browning, B.L., Zhou, Y. & Browning, S.R. A One-Penny Imputed Genome from Next-Generation Reference Panels. The American Journal of Human Genetics 103, 338–348 (2018).

60. Browning, S.R. & Browning, B.L. Rapid and Accurate Haplotype Phasing and Missing-Data Inference for Whole-Genome Association Studies By Use of Localized Haplotype Clustering. The American Journal of Human Genetics 81, 1084–1097 (2007).

61. McLachlan, G.J. Discriminant Analysis and Statistical Pattern Recognition. Wiley-Interscience (1992).

