## Supplementary Figures for "Polygenic Prediction of Complex Traits with Iterative Screen Regression Models"

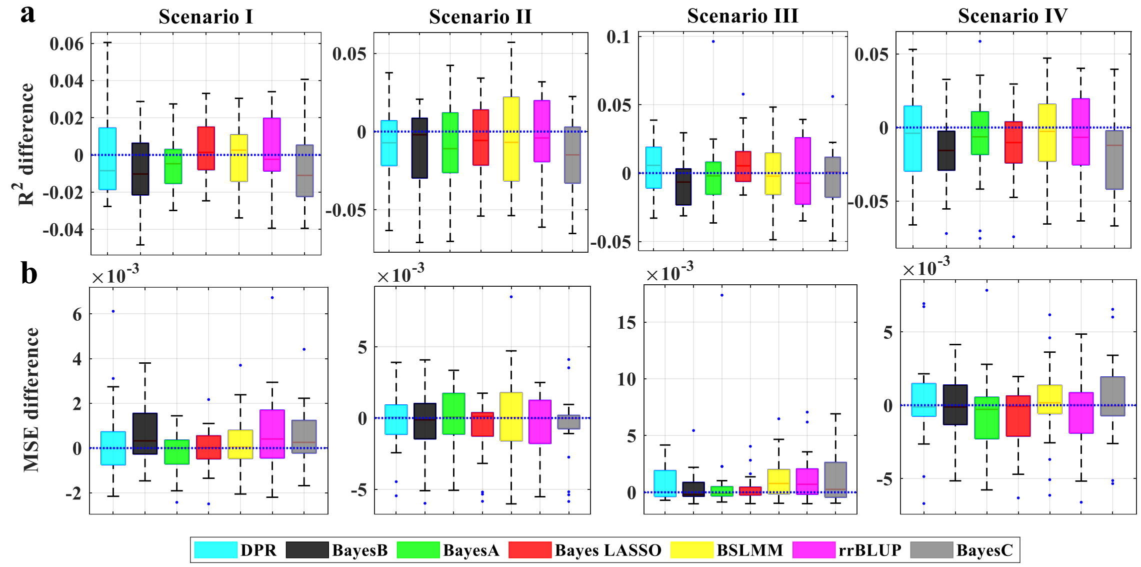


**Supplementary Figure 1. Comparison of prediction performance of seven methods with ISR in simulations when PVE = 0.2.** Performance is measured by $R^2$ difference (a) and MSE difference (b) with respect to ISR, where an $R^2$ difference below zero (i.e., values below the blue horizontal line) or a MSE difference above zero suggests worse performance than ISR. The sample R2 and MSE differences are obtained from 20 replicates in each scenario. Methods for comparison include DPR (cyan), BayesB (black), BayesA (green), Bayes LASSO (red), BSLMM (yellow), ), rrBLUP (purple), and BayesC (gray). Simulation scenarios include Scenario I, Scenario II, and Scenario III, which satisfies the DPR modeling assumption; where the number of SNPs in the large effect group is 100, 150, or 500; and Scenario IV, which satisfies the BayesR modeling assumption; For each box plot, the bottom and top of the box are the first and third quartiles, while the ends of whiskers represent either the lowest datum within 1.5 interquartile range of the lower quartile or the highest datum within 1.5 interquartile range of the upper quartile.

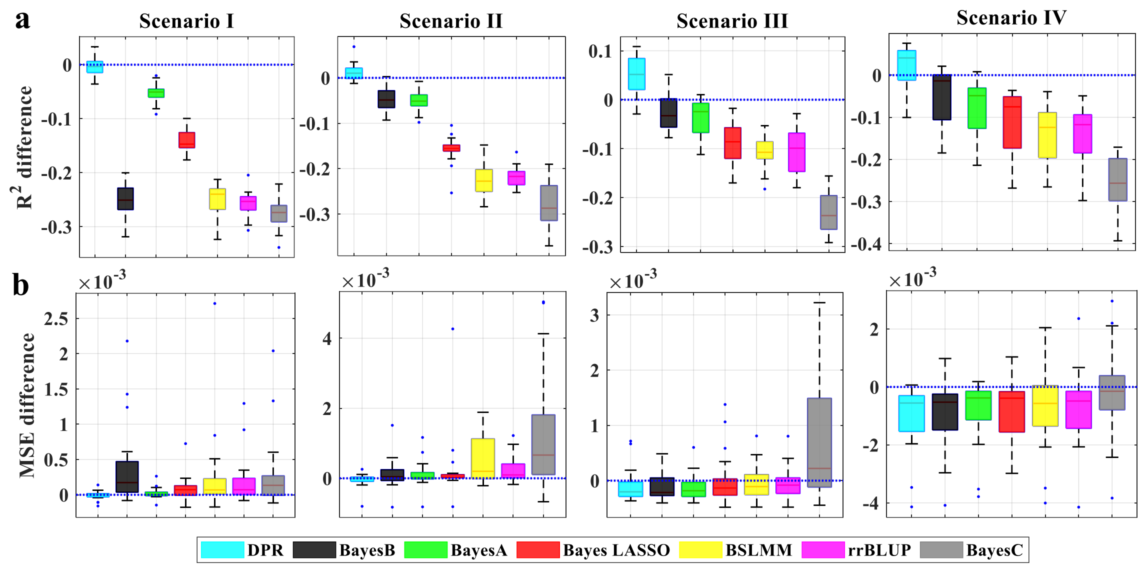


**Supplementary Figure 2. Comparison of prediction performance of seven methods with ISR in simulations when PVE = 0.8.** Performance is measured by $R^2$ difference (a) and MSE difference (b) with respect to ISR, where an $R^2$ difference below zero (i.e., values below the blue horizontal line) or a MSE difference above zero suggests worse performance than ISR. The sample R2 and MSE differences are obtained from 20 replicates in each scenario. Methods for comparison include DPR (cyan), BayesB (black), BayesA (green), Bayes LASSO (red), BSLMM (yellow), ), rrBLUP (purple), and BayesC (gray). Simulation scenarios include Scenario I, Scenario II, and Scenario III, which satisfies the DPR modeling assumption; where the number of SNPs in the large effect group is 100, 150, or 500; and Scenario IV, which satisfies the BayesR modeling assumption; For each box plot, the bottom and top of the box are the first and third quartiles, while the ends of whiskers represent either the lowest datum within 1.5 interquartile range of the lower quartile or the highest datum within 1.5 interquartile range of the upper quartile.

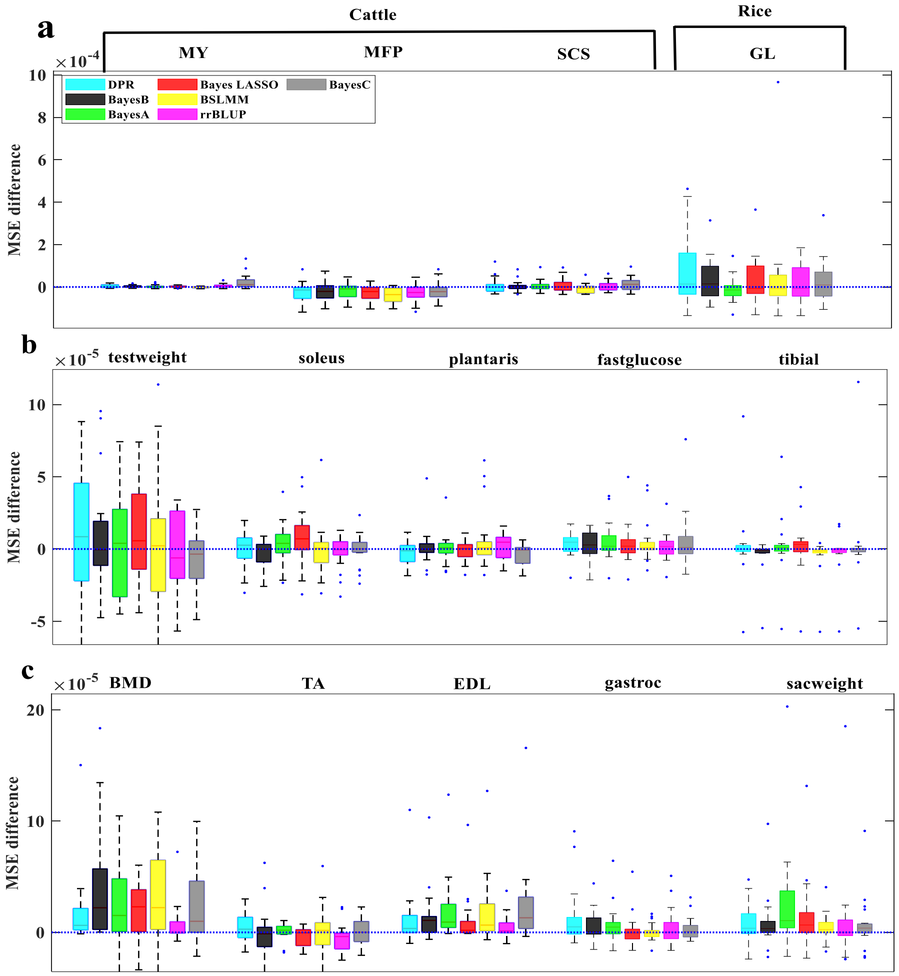


**Supplementary Figure 3. Comparison of prediction performance of several methods with DPR.MCMC for twelve traits from three data sets.** a Prediction performance for MFP, MY, and SCS in the cattle data, and for GL in the rice data; b,c Prediction performance for the ten traits in the mice data. Performance is measured by MSE difference with respect to ISR, where an MSE difference above zero suggests worse performance than ISR. The sample MSE differences are obtained from 20 replicates in each scenario. Methods for comparison include DPR (cyan), BayesB (black), BayesA (green), Bayes LASSO (red), BSLMM (yellow), ), rrBLUP (purple), and BayesC (gray). For each box plot, the bottom and top of the box are the first and third quartiles, while the ends of whiskers represent either the lowest datum within 1.5 interquartile range of the lower quartile or the highest datum within 1.5 interquartile range of the upper quartile.

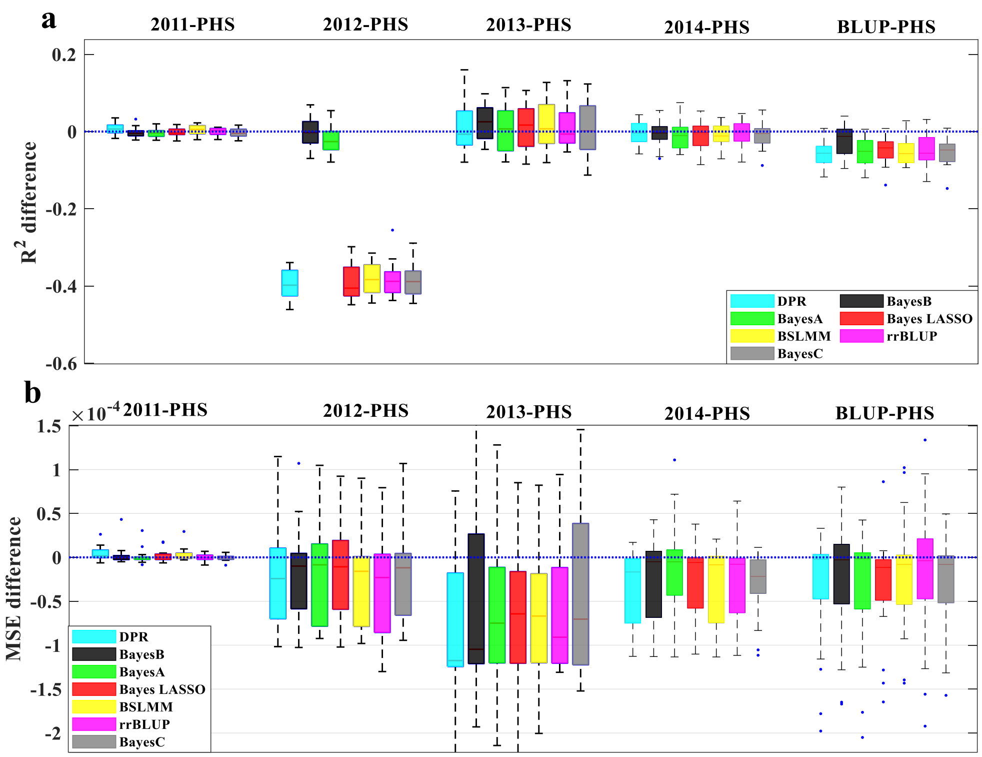


**Supplementary Figure 4. Comparison of prediction performance of seven methods with ISR for fourteen traits from three data sets.** Prediction performance for PHS with difference years in the wheat data; a,b Performance is measured by R2 difference and MSE difference with respect to ISR, where a negative value (i.e., values below the red horizontal line) or above zero suggests worse performance than ISR, respectively. Methods for comparison include DPR (cyan), BayesB (black), BayesA (green), Bayes LASSO (red), BSLMM (yellow), rrBLUP (purple), and BayesC (gray). For each box plot, the bottom and top of the box are the first and third quartiles. At the same time, the ends of whiskers represent either the lowest datum within 1.5 interquartile range of the lower quartile or the highest datum within 1.5 interquartile range of the upper quartile. The sample R2 differences and MSE difference are obtained from 10 fold validation for each trait.

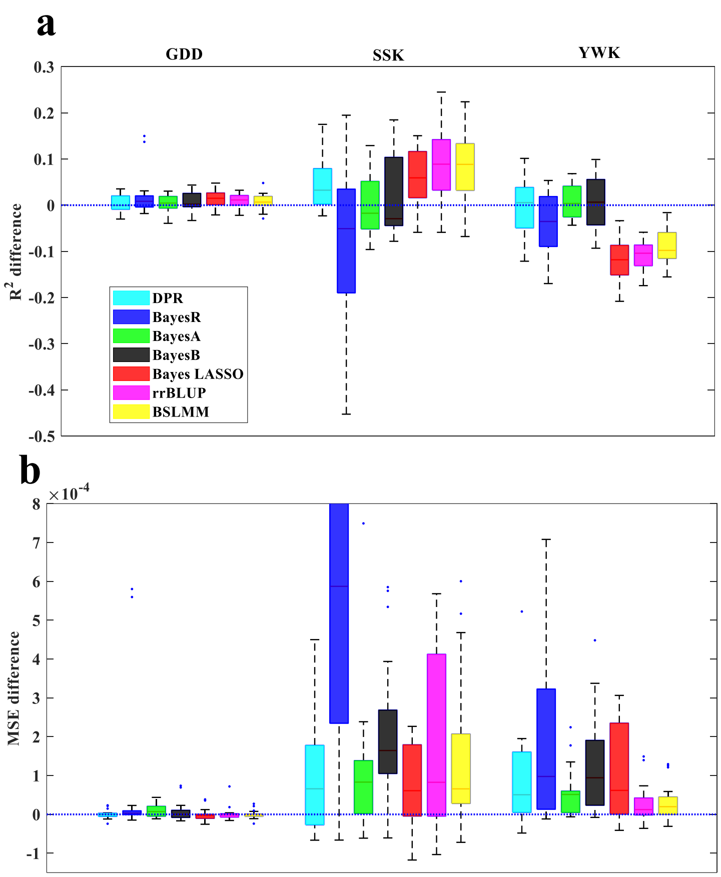


**Supplementary Figure 5. Comparison of prediction performance of seven methods with ISR for three traits from maize data sets.** a,b Performance is measured by R2 difference and MSE difference with respect to ISR, where a negative value (i.e., values below the red horizontal line) or above zero suggests worse performance than ISR, respectively. Methods for comparison include DPR (cyan), BayesR(Blue), BayesA (green), BayesB (black), Bayes LASSO (red), rrBLUP (purple), BSLMM (yellow). For each box plot, the bottom and top of the box are the first and third quartiles, while the ends of whiskers represent either the lowest datum within 1.5 interquartile range of the lower quartile or the highest datum within 1.5 interquartile range of the upper quartile.

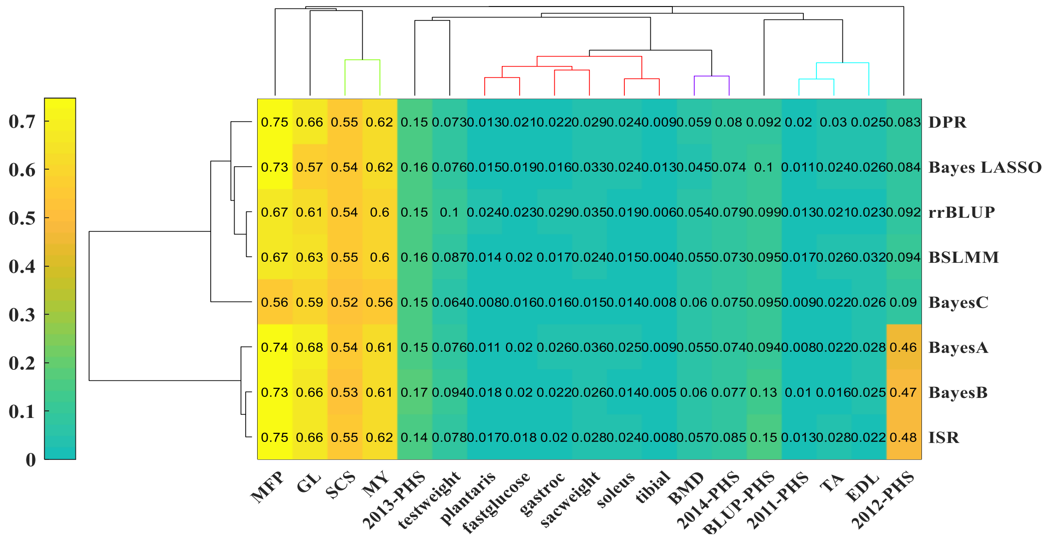


**Supplementary Figure 6. The clustering result with heatmap.** Based on the Simulations and Real data applications results the averaged prediction of $R^2$

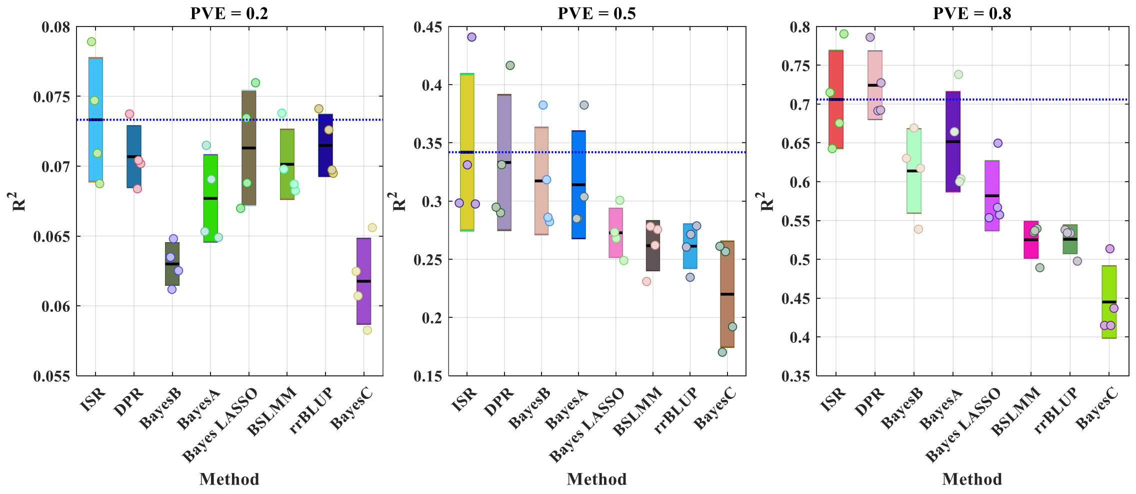


**Supplementary Figure 7. The clustering result with heatmap.** Based on the simulations data applications results in the averaged prediction of $R^2$. Box plots 95% SEM and SD as a box using patch objects. Here, the SD equals the mean above the patch. The tash line was the means predictive of ISR $R^2$.

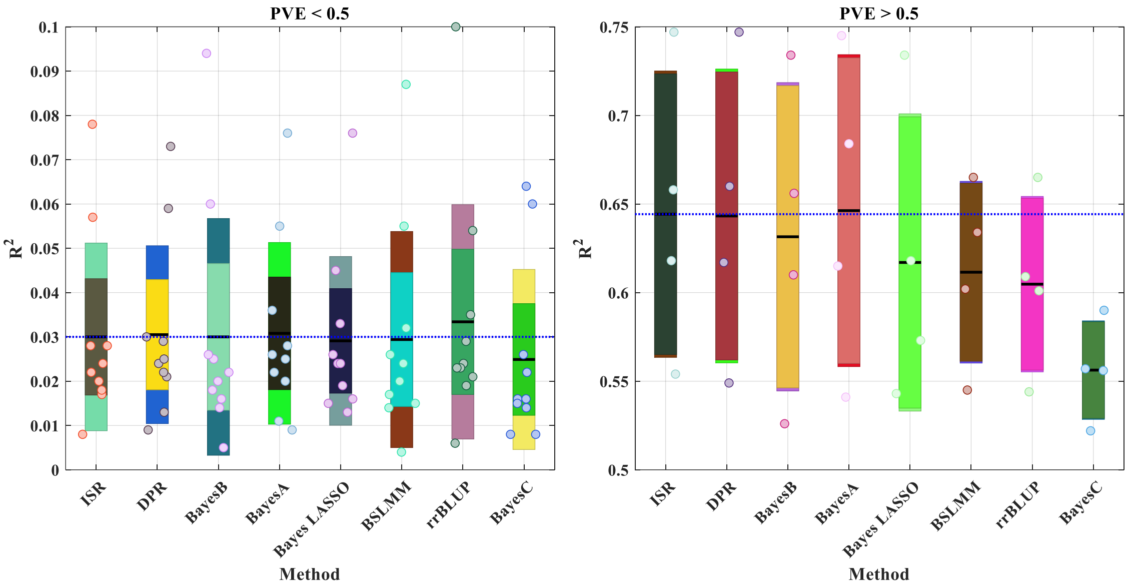


**Supplementary Figure 8. The clustering result with heatmap.** Based on the Real data applications results the averaged prediction of $R^2$

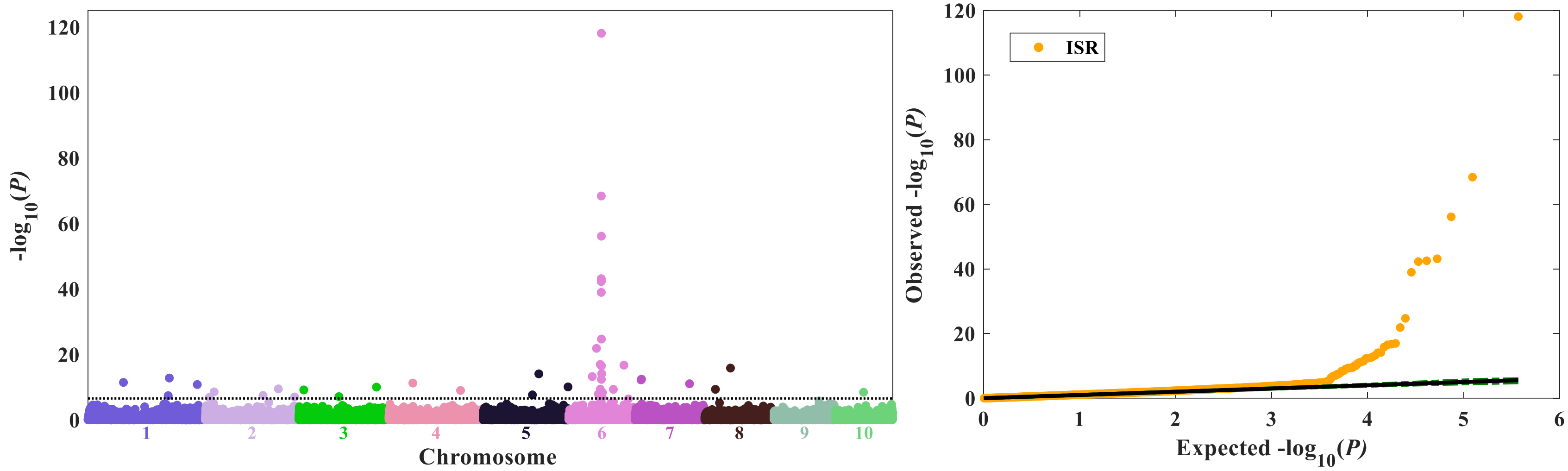


**Supplementary Figure 9. Genome-wide association study (GWAS) for yellow versus white kernels.** GWAS for kernel color on 1,595 maize inbred lines with yellow or white kernels. The black dash line in the Manhattan plot shows that the value of Bonferroni multiple test correction (0.05/p, p is the number of SNPs), and the green dash lines in the QQ plot show that the 95% confidence intervals.

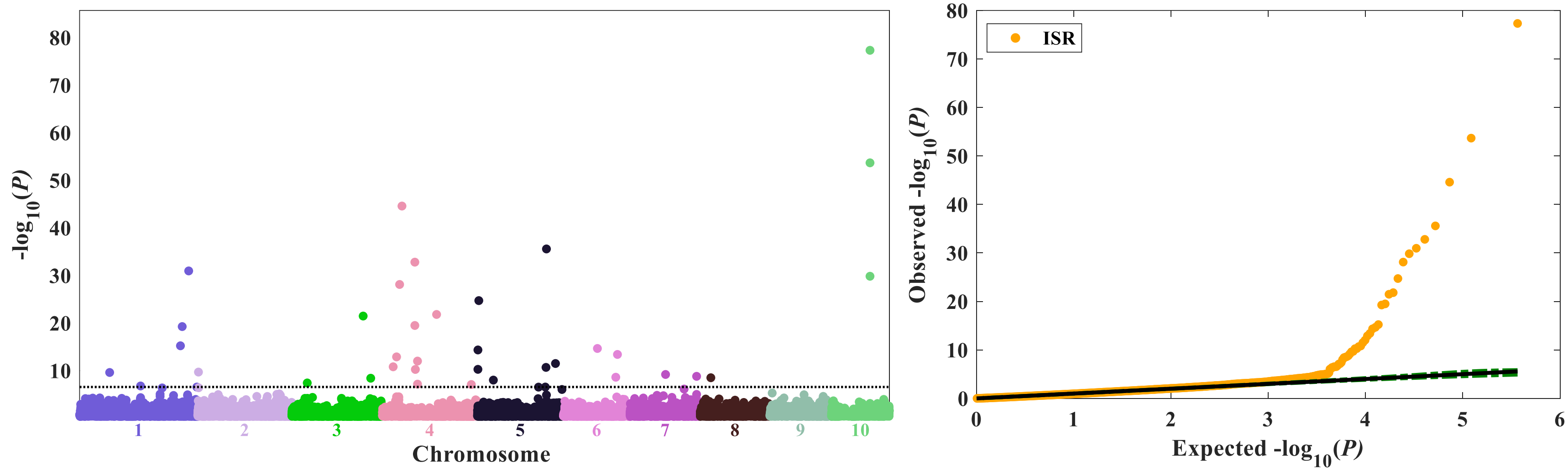


**Supplementary Figure 10. Genome-wide association study (GWAS) for sweet versus starchy corn.** GWAS for kernel color on 2,145 maize inbred lines with sweet or starchy kernels. SNP, single-nucleotide polymorphism. The black dash line in the Manhattan plot shows that the value of Bonferroni multiple test correction (0.05/p, p is the number of SNPs), and the green dash lines in the QQ plot show that the 95% confidence intervals.

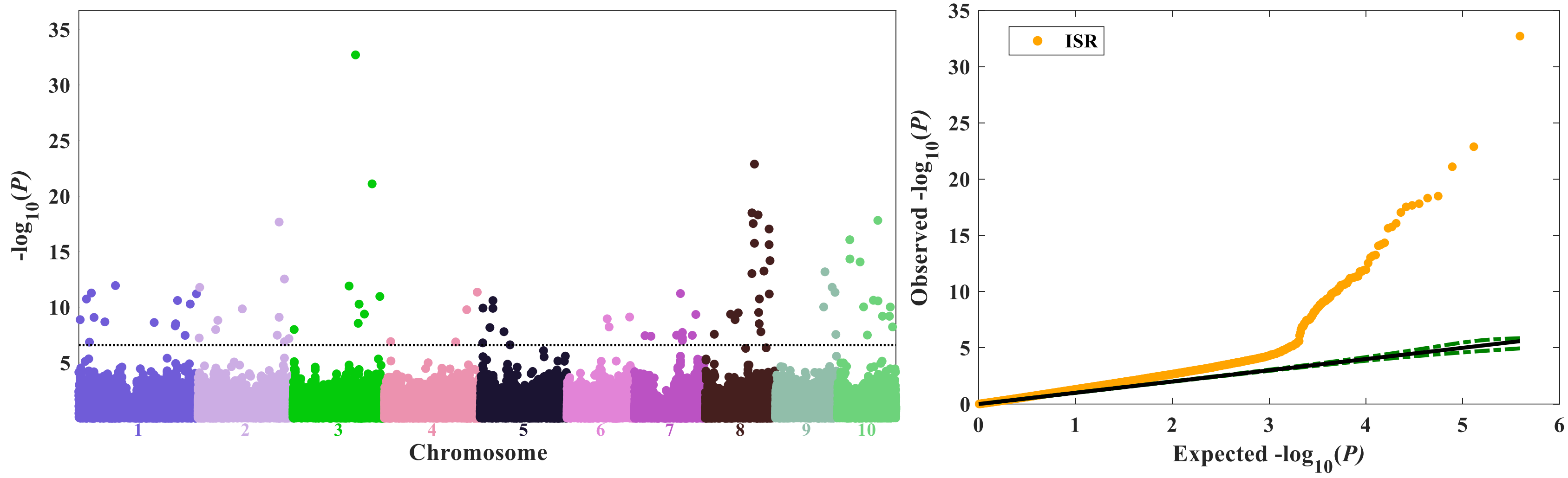


**Supplementary Figure 11. Genome-wide association study (GWAS) for growing degree days to silking.** GWAS for growing degree days to 50% silking on 2,279 maize inbred lines.The black dash line in the Manhattan plot shows that the value of Bonferroni multiple test correction (0.05/p, p is the number of SNPs), and the green dash lines in the QQ plot show that the 95% confidence intervals.
